# Molecular docking, validation, dynamic simulation and pharmacokinetic prediction of natural compounds against *Mycobacterium tuberculosis* Beta-lactamase BlaC

**DOI:** 10.1101/2025.01.07.631633

**Authors:** Sandra Sreekumar, J.L Bhagya Jyothi, S.S Liji, Subhash Chandra Pani, Niya Elizaba Rajee, Aguero Pizzolo Stephanie, Terreux Raphael, Immanuel Dhanasingh

## Abstract

Tuberculosis, caused by *Mycobacterium tuberculosis* ranks second globally in terms of infectious disease-related deaths, after HIV. The resistance of *M. tuberculosis* to existing beta-lactam antibiotics is primarily due to chromosomally encoded gene blaC, which can hydrolyse predominantly available beta-lactam antibiotics. Despite the available beta-lactamase inhibitors with beta-lactam ring such as clavulanate, being efficacious, they lead the bacteria to develop an inhibitor escape mechanism. In contrast, the natural product inhibitors without beta-lactam ring that might resist bacterial escape mechanisms have the additional advantage of fewer side effects, higher bioavailability, and better bioremediation towards environmental sustainability. This study identifies novel natural inhibitors against BlaC, from the Natural Products Atlas database of PubChem and ZINC comprising 10,000 compounds using molecular docking and Molecular Dynamics (MD) simulations. The virtual screening and docking experiments demonstrated that the top 10 compounds exhibited favourable docking scores in the range of -7.59 to -9.63 kcal/mol compared to the reference molecule, Doripenem (-7.35 kcal/mol). Following targeted docking and ADMET analysis, three lead compounds Lecanorafuran A, Tryptoquivaline K, and Deacetylisowortmin A were selected for MD simulations for a period of 100 ns to evaluate the stability of the protein-ligand complexes. Based on Root Mean Square Deviation, Root Mean Square Fluctuation, Radius of gyration and Solvent Accessible Surface Area, PCA, and FEL analysis, it was found that Tryptoquivaline K exhibited consistent stability throughout the simulation. Additionally, Molecular Mechanics/Poisson-Boltzmann Surface Area (MMPBSA) analysis was conducted to assess the binding affinity of the complex. All these analyses validate that the natural compound Tryptoquivaline K exhibits potential as a promising BlaC inhibitor but warrants further validation through experimental studies in this aspect.

## 1. INTRODUCTION

Tuberculosis (TB) is an extremely infectious disease caused by the gram-positive bacterium, *Mycobacterium tuberculosis* that most often affects the lungs and other parts of the body. It spreads through the air when TB-infected patients cough, sneeze, and spit [1]. The primary symptoms of TB include persistent cough for more than 2 weeks, weight loss, fever, chest pain, and fatigue [2]. Despite significant advancements in diagnosis and treatment over the decades, the disease continues to rank second globally in terms of infectious disease-related deaths, after HIV [3]. This is due to the following reasons:1) lack of vaccines for adults, 2) its long doubling time, 3) intracellular localisation in macrophages, 4) the presence of mycolic acid in the bacterial cell wall that creates a hydrophobic barrier against antibiotics, and 5) the rise of novel drug-resistant strains. According to the World Tuberculosis Report, 10.6 million individuals worldwide were affected by tuberculosis in 2022, equivalent to 133 cases per 1,00,000 people. A total of 1.3 million TB-related deaths worldwide and 1,67,000 HIV-related fatalities have been reported. In 2022, eight countries accounted for more than two-thirds of global TB cases: India (27%), Indonesia (10%), China (7.1%), Philippines (7%), Pakistan (5.7%), Nigeria (4.5%), Bangladesh (3.6%) and the Democratic Republic of Congo (3%) (*1.1 TB Incidence*, n.d.). Consequently, tuberculosis caused by the gram-positive intracellular bacterium *M. tuberculosis* presents a considerable risk of morbidity to humans in developed and developing areas, making it a serious global health concern (Bloom et al., 2017) (*Tuberculosis*, n.d.).

The medications used to treat TB are divided into two categories based on the present therapeutic regimen: first-line and second-line (World Health Organization, 2014). Rifampicin, isoniazid, pyrazinamide, and ethambutol are FDA-approved first-line medications for *Mycobacterium tuberculosis* infections, while, the second line of treatment consists of injectable medications like capreomycin, kanamycin, amikacin, fluoroquinolones such as levofloxacin, moxifloxacin, and gatifloxacin (Seung et al., 2015). Further, certain combinations of beta-lactam antibiotics involving anti-TB-drugs such as meropenem, bedaquiline, and amoxicillin along with beta-lactam inhibitors such as clavulanate, are also being routinely used in MDR-TB treatment (Zhang et al., 2015). Despite these treatment strategies, there is an alarming increase in the number of multi-drug resistant *M.tuberculosis* strains (Seung et al., 2015). Multi-drug resistant tuberculosis or MDR-TB, one of the complications of tuberculosis therapy which was identified in the late 1980s and early 1990s characteristically shows resistance to first-line medications such as rifampicin, second-line aminoglycoside, and to fluoroquinolones (Seung et al., 2015). For instance, an extensive report from WHO estimated that 4,10,000 people were affected by Rifampicin-resistant TB (RR-TB) and an estimated 1,91,000 deaths occurred due to MDR-TB/RR-TB in 2021 (*2.3 Drug-Resistant TB*, n.d.). Thus, the urgent need for innovative drug candidates is critical for effective treatment strategies, as multidrug-resistant (MDR) and extensively drug-resistant (XDR) strains have made tuberculosis therapy increasingly vital.

The resistance to beta-lactam antibiotics in *M. tuberculosis* is primarily due to the presence of chromosomally encoded gene *blaC,* whose product beta-lactamase enzyme is capable of hydrolyzing beta-lactam antibiotics that are currently in use against TB (Hugonnet & Blanchard, 2007). BlaC (28.4 kDa, 307 amino acids), the monomeric β-lactamase of *M. tuberculosis*, belongs to Ambler class-A that exhibits a wide range of substrate specificity and is capable of hydrolyzing various β-lactam antibiotics, including penicillins, cephalosporins and carbapenems such as imipenem, which complicates the use of these drugs in treating tuberculosis (Wang et al., 2006). Therefore, the BlaC enzyme has already been validated as one of the leading therapeutic targets against *M. tuberculosis* (Clement et al., 2021).

The development of potential inhibitor molecules against BlaC has been warranted for the effective use of beta-lactam antibiotics against tuberculosis. Prior studies have identified that the β-lactamase inhibitor, clavulanate from *Streptomyces clavuligerus* could slowly, but irreversibly inhibit BlaC (Hugonnet & Blanchard, 2007). This accelerated the development of a treatment combining β-lactam antibiotic and clavulanate against MDR/XDR TB. When combined with traditional β-lactams such as amoxicillin and ticarcillin, clavulanic acid produces a broad spectrum of antibiotic combinations effective against many β-lactamases producing bacteria (Shah et al., 2024). A conjugate of meropenem with clavulanate was developed and found to be effective against XDR strains of *M. tuberculosis* (Hugonnet et al., 2009). However, it was found that a point mutation N132G in BlaC enables it to efficiently hydrolyse beta-lactam containing clavulanate and therefore, completely escape the inactivation [22]. As a result, alternative treatment strategies, such as identifying non-beta lactam inhibitors are often necessary to combat *M. tuberculosis* effectively.

Natural products have been widely recognized as the foundation for new therapeutic lead compounds due to their fewer side effects, and higher bioavailability following their vast distribution among plants, animals, marine life, and microorganisms (Mushtaq et al., 2018). Many natural compounds are currently being examined through computational approaches for their bioactivity with the aim of recent developments in proteomics, genomics, and transcriptomics, which may aid in developing new drugs in the future (Mushtaq et al., 2018; Shah et al., 2024). Unlike chemical compounds, natural products can be easily bioremediated to maintain environmental sustainability. Therefore, the identification and discovery of natural compound inhibitors become a priority in the fight against TB.

Here, we have applied computational approaches to identify natural inhibitors against the M. tuberculosis BlaC beta-lactamase enzyme. In the current study, 10,000 natural compounds from the Natural Products Atlas database of PubChem (Kim et al., 2019; Li et al., 2010) and ZINC (Mathpal et al., 2022) databases were screened against the β-lactamase (BlaC) of *Mycobacterium tuberculosis* to identify a novel therapeutic drug candidate that can be utilized for effective beta-lactamase inhibition, which can further serve as an imperative contribution in the effective tuberculosis treatment against MDR/XDR variants.

## 2. MATERIALS AND METHODS

### 2.1 Receptor preparation

The resolved co-crystal structure of *M. tuberculosis* β-lactamase (BlaC) with Doripenem (DRW) having a resolution of 2.20 Å (PDB ID:3IQA) was retrieved from the Protein Data Bank (https://www.rcsb.org) (Fig. 1a, b) (Zardecki et al., 2016). For protein preparation, all water molecules and ions were removed from the protein molecule using PyMOL (version 3.1) software (Yuan et al., 2017). Further, the addition of hydrogen atoms to the receptor molecule was carried out by using the Autodock 4.2.6 software [23]. The structure of the protein was saved in .pdb format for further analysis.

**Fig. 1.**
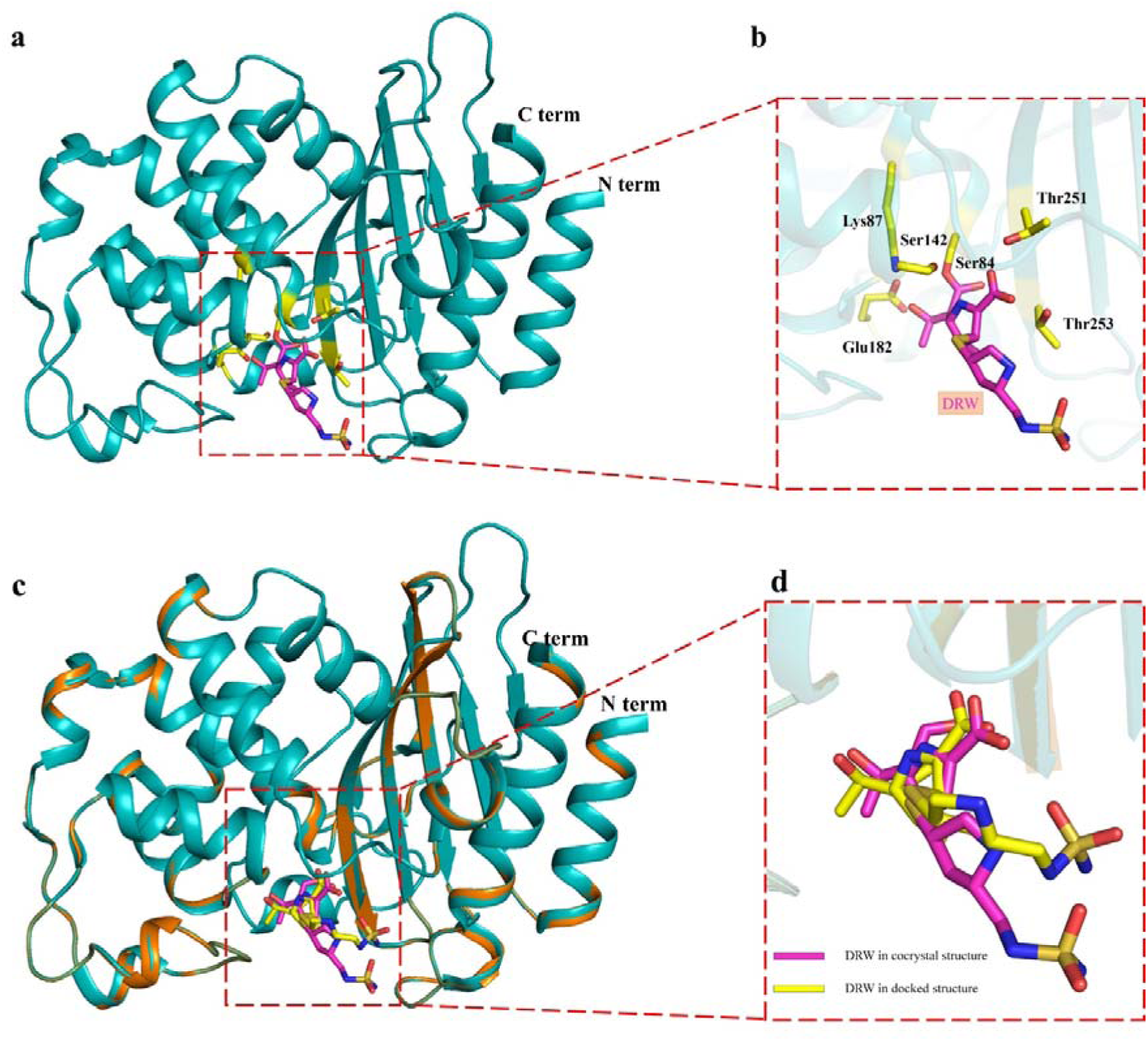
**a** Crystal structure of *Mycobacterium tuberculosis* β-lactamase BlaC (PDB ID: 3IQA) represented in cartoon model with a red dotted box showing the active site consisting of ligand doripenem represented in magenta stick model **b** Close-up view of the active site with its critical residues represented as yellow sticks and the ligand doripenem in magenta colour **c** Superimposition of the crystal structure of *Mycobacterium tuberculosis* β-lactamase BlaC (PDB ID: 3IQA) represented as cartoon in Cyan with the docked structure of BlaC (represented as cartoon in orange) **d** Close-up view of the superimposed ligands; Doripenem of reference BlaC and doripenem of docked BlaC in magenta and yellow stick models respectively

### 2.2 Ligand Preparation

A library of 10,000 compounds was retrieved from ZINC and Natural Products Atlas of PubChem databases in .sdf format. Open Babel (version 3.1.0) (O’Boyle et al., 2011) converted the .sdf file to .pdb format. A natural substrate of BlaC, Doripenem (DRW) or (2S,3R,4S)-2-[(2S,3R)-3-hydroxy-1-oxobutan-2-yl]-3-methyl-4-({(3S,5S)-5-[(sulfamoylamino)methyl]pyrrolidine-3-yl}sulfanyl)-3,4-dihydro-2H-pyrrole-5-carboxylic acid was taken as a reference molecule, whose structure was downloaded from PubChem database (https://pubchem.ncbi.nlm.nih.gov) for comparative analysis as a control (Kim et al., 2019). Ligands were prepared by adding hydrogens to all the compounds. Energy minimization was done with the Universal Force Field (UFF) (Artemova et al., 2016) using a conjugate-gradient algorithm in PyRx software (Mun et al., 2022). The resulting compounds were saved in .pdbqt format.

### 2.3 Structure-Based Virtual Screening

To find potential compounds against *Mycobacterium tuberculosis* β-lactamase (BlaC), the Autodock Vina module in PyRx software (GUI version 0.8) was used to screen conformations of ligands with high affinity at the active site of beta-lactamase using molecular docking (Trott & Olson, 2010). Virtual screening was performed for the ligands collected from the ZINC and Natural Products Atlas of PubChem database against the target *Mycobacterium tuberculosis* β-lactamase. The protein and ligand structures prepared in the previous step were used for virtual screening.

### 2.4 Docking Validation

Initially, molecular docking analysis was performed with the reference molecule doripenem (DRW) in the active site of beta-lactamase to re-produce the same conformation similar to the co-crystallized ligand. The grid centre for docking was set as X = -7.765037, Y = -8.462741, and Z= 2.931704 Å with dimensions of the grid box 40 x 40 x 40 Å. Re-docking using Autodock 4.2.6 was also performed to validate the docking accuracy of the software PyRx. The top 10 compounds selected from virtual screening based on binding energy were employed for docking using the Autodock program. The grid center was set with the same XYZ coordinates as that of the reference docking. Finally, the protein-ligand complex with the lower binding energy as compared to the reference molecule was selected for further molecular dynamic simulation.

### 2.5 Visualization

The 2D interactions of protein-ligand complexes including hydrogen bonds and the bond lengths were analysed using Discovery Studio Visualiser software (version 3.5). At the same time, further scrutiny of the 3D structure for stable interactions was visualized using the PyMOL (version 3.1) molecular visualization tool.

### 2.6 Pharmacokinetic properties and Lipinski’s rule of five

Pharmacokinetic properties are essential in determining drug molecule’s absorption, distribution, metabolism, and excretion characteristics in the human body. Ligands with higher binding energy values against BlaC were selected for evaluation via insilico ADMET analysis, to predict their pharmacokinetic analysis, drug likeliness, and toxicity analysis.

ADMET analysis of the top hits including Lipinski’s rule of five were examined. Lipinski’s oral drug likeliness properties include i) molecular weight (<500 Daltons), ii) Number of hydrogen bond donors (<5), iii) number of hydrogen bond acceptors (<10), and iv) Log P (<5) (Lipinski et al., 2001). The ADMET properties such as P-glycoprotein inhibitor/substrate, human intestinal absorption (HIA), blood-brain barrier penetration, and AMES toxicity were investigated (Aier, n.d.). The prediction was carried out using ProTox (https://tox.charite.de) and SwissADME (https://www.swissadme.ch/).

### 2.7 Molecular Dynamic Simulations

The MD Simulation is widely used to assess the structural stability of the protein-ligand complexes under physiological conditions (Hollingsworth & Dror, 2018). The selected compounds were redocked against BlaC with Autodock 4.2.6 using the same docking protocol, repeating five times for each compound. The docking pose was strictly scrutinized using PyMOL (version 3.1) for stable interactions. The complexes with the best pose that had maximal interaction with the active residues were chosen for MD simulations. All MD Simulations were performed using the Gromacs 2022.4.1 software package, and the complexes’ topologies were prepared using the CHARMM 36 force field (Bjelkmar et al., 2010). Solvation was achieved using the TIP3P water model (Mark & Nilsson, 2001). After solvation, neutralization of the system was done by the addition of 44 Na^+^ and 38 Cl^-^ ions. Further, energy minimization was carried out to ensure that the system has no steric clashes. This step was done with the steepest descent algorithm by using the Verlet cut-off scheme (Diem & Oostenbrink, 2020). After energy minimization, equilibration was conducted under the NVT method at 300 K for 10 ps. The next phase of equilibration was carried out under the NPT method at 1 atm for 10 ps. The system was subjected to a constant temperature of 300 K and a constant pressure of 1 atm with a time step of 2 fs using Parrinello-Rahman for constant pressure simulations. All MD simulations were conducted for 100 ns and the stability of the control and the complexes were calculated by analyzing the Root Mean Square Deviation (RMSD), Root Mean Square Fluctuation (RMSF), Radius of gyration (Rg), Solvent Accessible Surface Area (SASA), Principal Component Analysis and Free Energy Landscape analysis. This analysis helps to examine different structural and dynamic characteristics of a simulated system, including flexibility and interactions with surrounding solvents.

### 2.8 Binding free energy analysis

The binding free energy analysis reveals the stability of the ligand in the active site of the protein through various interactions (Mathpal et al., 2022). A detailed analysis of the binding free energy was performed using the Molecular Mechanics Poisson–Boltzmann Surface Area (MM-PBSA) protocol implemented in the g_mmpbsa package. MM-PBSA calculation quantitatively estimates the interaction mechanisms of the proteins and the ligand molecules (Mathpal et al., 2022).

## 3. RESULTS AND DISCUSSIONS

### 3.1 Virtual Screening

The co-crystal structure of *M. tuberculosis* β-lactamase (BlaC) (PDB ID: 3IQA) was retrieved from the RCSB Protein data bank. The crystal structure had a completely built model between amino acids D43 and V307 excluding the signal peptide (M1-G35). The model was prepared as mentioned above and then subjected to multiple ligand docking using PyRx software. Virtual screening was performed for the 10,000 compounds of the Natural Products Atlas (NPA) database of PubChem and ZINC database. The top 10 compounds (Table. 1) with binding energy ranging between -9.5 to -10.6 kcal/mol, which was lower or equivalent to the binding energy of the reference molecule Doripenem (DRW) (-7.35 kcal/mol) were selected for further studies.

**Table 1.** Global docking using PyRx results of the top 10 natural compounds from the NPAtlas from database of PubChem.

| Sl. No | Compound ID | Compound Name | Binding energy(kcal/mol) |
| --- | --- | --- | --- |
| 1 | NPA013206 | Aspergilazine A | -10.6 |
| 2 | NPA000175 | Aschernaphthopyrone A | -10.4 |
| 3 | NPA013325 | Naseseazine A | -10.4 |
| 4 | NPA000079 | 25-methylpisolactone | -10.3 |
| 5 | NPA008469 | (-)-R-skyrin | -10.2 |
| 6 | NPA000475 | Chrondramide 7c | -10.2 |
| 7 | NPA001729 | Lecanorafuran A | -10.1 |
| 8 | NPA011913 | Tryptoquivaline K | -9.9 |
| 9 | NPA002942 | Deacetylisowortmin A | -9.9 |
| 10 | NPA011424 | Chaiyaphumine B | -9.7 |

### 3.2 Docking Validation

To validate the docking parameters in Autodock 4.2.6, the prepared protein molecule was subjected to docking with the known, co-crystallized ligand Doripenem (Fig. 1c). The docked pose was compared with the co-crystal structure (PDB: 3IQA) (Fig. 1c). Upon superimposition of the docked model with co-crystal structure, it aligned with an RMSD value of 0.00 Å (Fig. 1c, d). The active site residues of BlaC protein comprises of the catalytic Ser70, and the surrounding Lys73, Ile117, Ser130, Glu166 and Thr251 are involved in the hydrolysis of the substrate (Tremblay et al., 2010). Analysis of the interaction between β-lactamase (BlaC) and doripenem (DRW) using Discovery Studio Visualizer software indicated that six hydrogen bonds were formed with the active site residues, namely, Ser84, Lys87, Ser142, Glu182, Thr251, and Thr253 (Fig. 2a). While the *insilico* docked BlaC-DRW exhibited five hydrogen bonds with active residues Ser84, Lys87, Ser142, Thr253 and Glu292 (Fig. 2b). The protein-ligand interaction analysis revealed that the docked complex forms a similar pose, and interactions compared to the crystal structure, strongly validating the docking protocol. Furthermore, based on the validated docking protocol and identified active site residues, the top ten compounds obtained from virtual screening (Table. 2) were docked against BlaC using Autodock 4.2.6. The evaluated binding energy of the 10 compounds is given in Table 2. Docking results revealed that all 10 compounds had lower binding energy in the range of -7.59 to -9.63 kcal/mol than the reference molecule (DRW) (-7.35 kcal/mol), which further substantiates the virtual screening results. Thus, the docking analysis confirmed the strong binding affinity of the top-hit compounds with the target protein, suggesting a more detailed investigation to understand their characteristics.

**Fig. 2.**
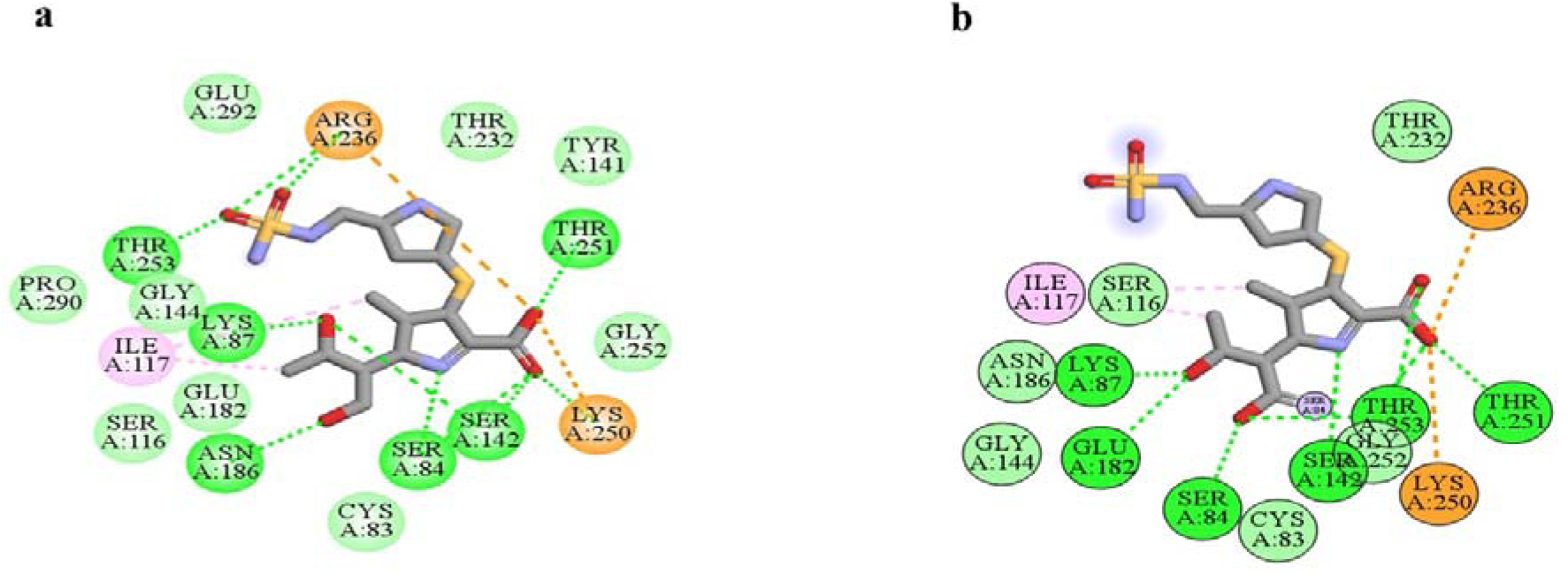
**a** 2 D ligand interaction of X-ray crystal structure of BlaC with DRW (PDB ID:3IQA) **b** 2 D interaction of docked complex DRW with BlaC (Dotted green lines indicate the hydrogen bond and orange dotted lines represent pi-interaction)

**Table 2.**
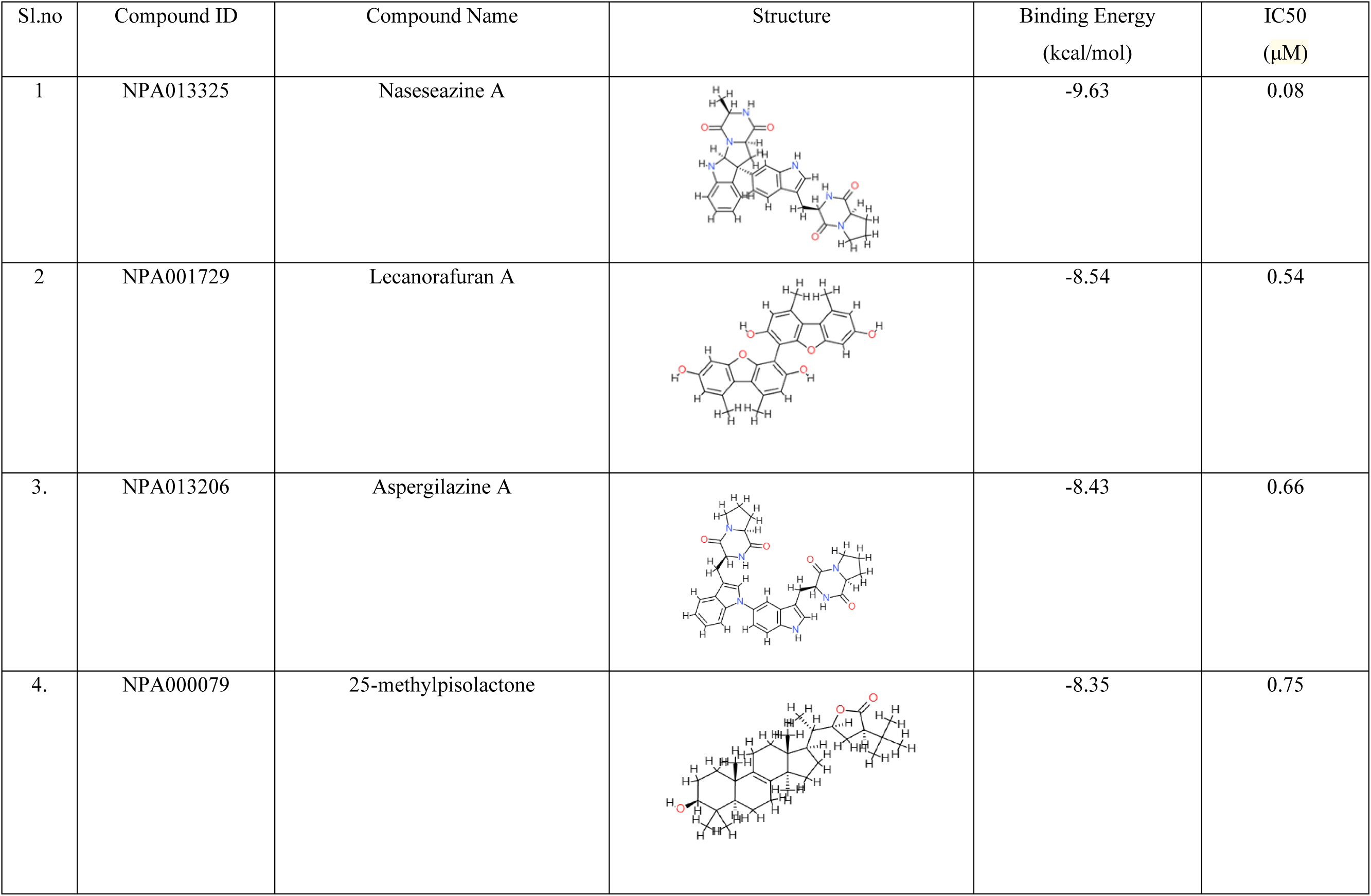
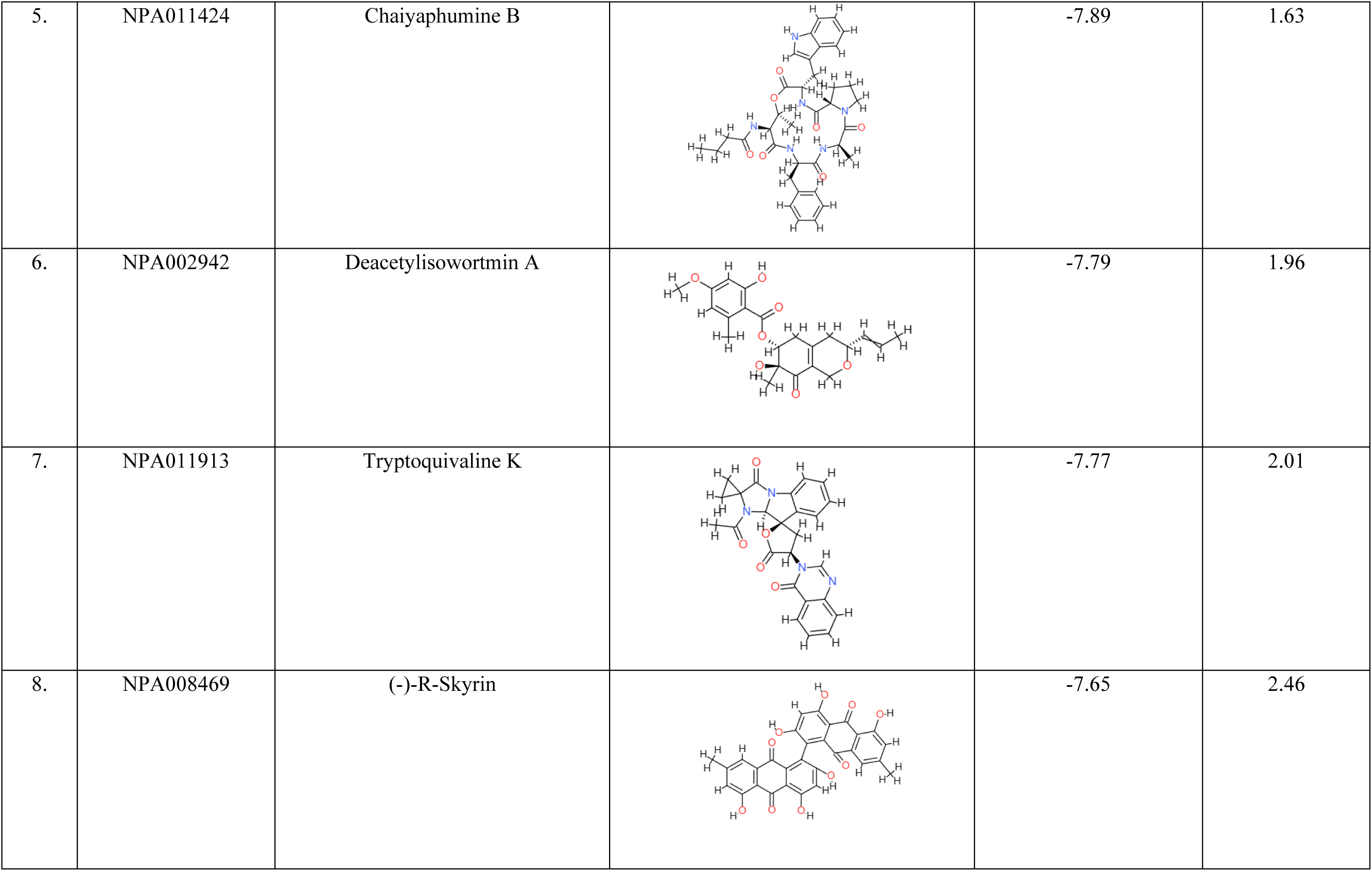
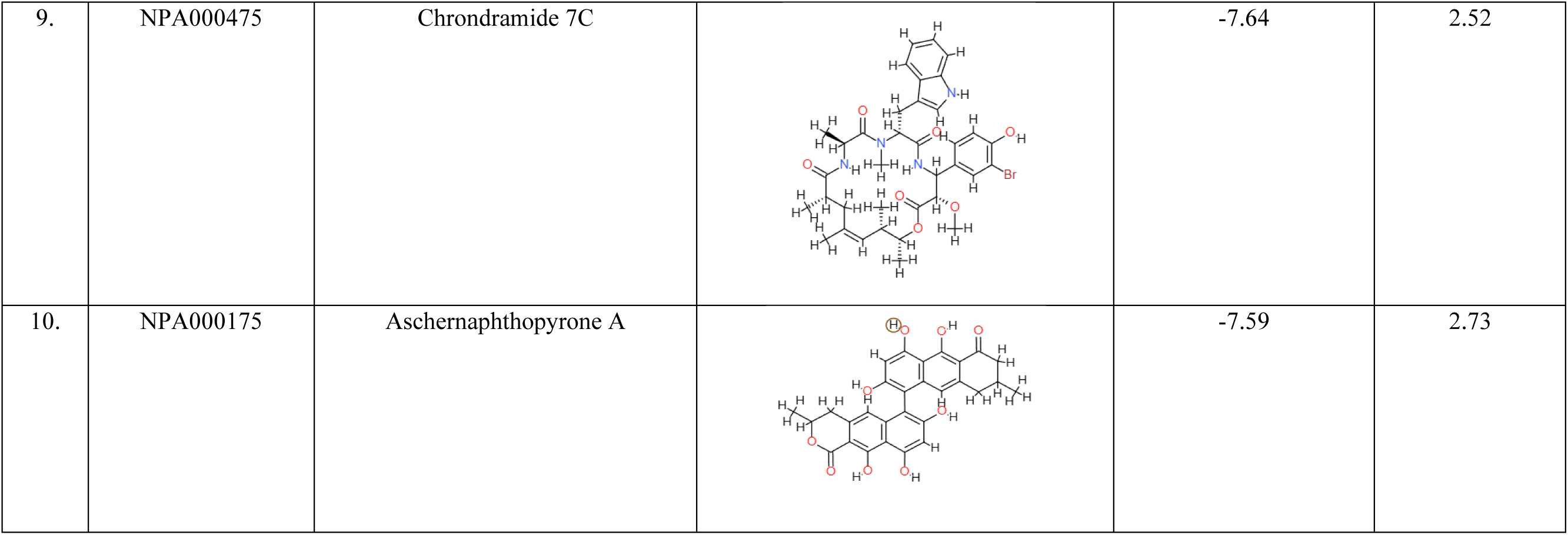
Binding energy of the top 10 compounds selected for site-specific docking using AutoDock 4.2.

### 3.3 Molecular Interactions of top ligands with BlaC

The 2D interactions of the top 10 hits and the reference compound were visualized using Discovery Studio Visualizer software (Table 3 and Fig. 3). The docked poses demonstrated that the compounds bind within the active site of the *M. tuberculosis* β-lactamase, BlaC, forming stable interactions with the surrounding residues.

**Fig. 3.**
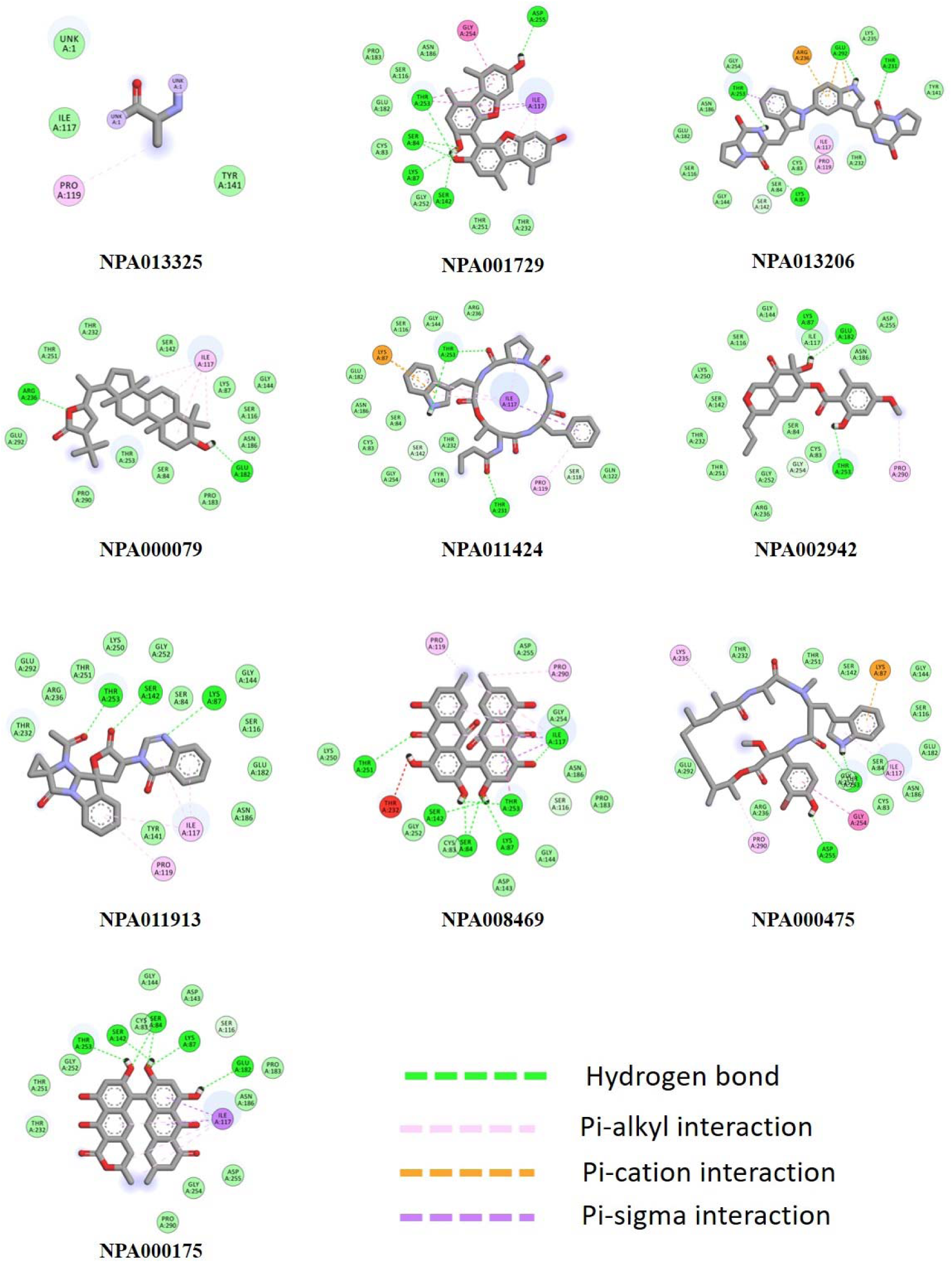
2 D protein-ligand interaction diagrams of top10 ligands with BlaC (In all compounds, dotted, dark green lines indicate the hydrogen bond, lavender lines indicate pi-alkyl interactions, orange line shows pi-cation interactions, purple line specifies pi-sigma interactions

**Table 3.**
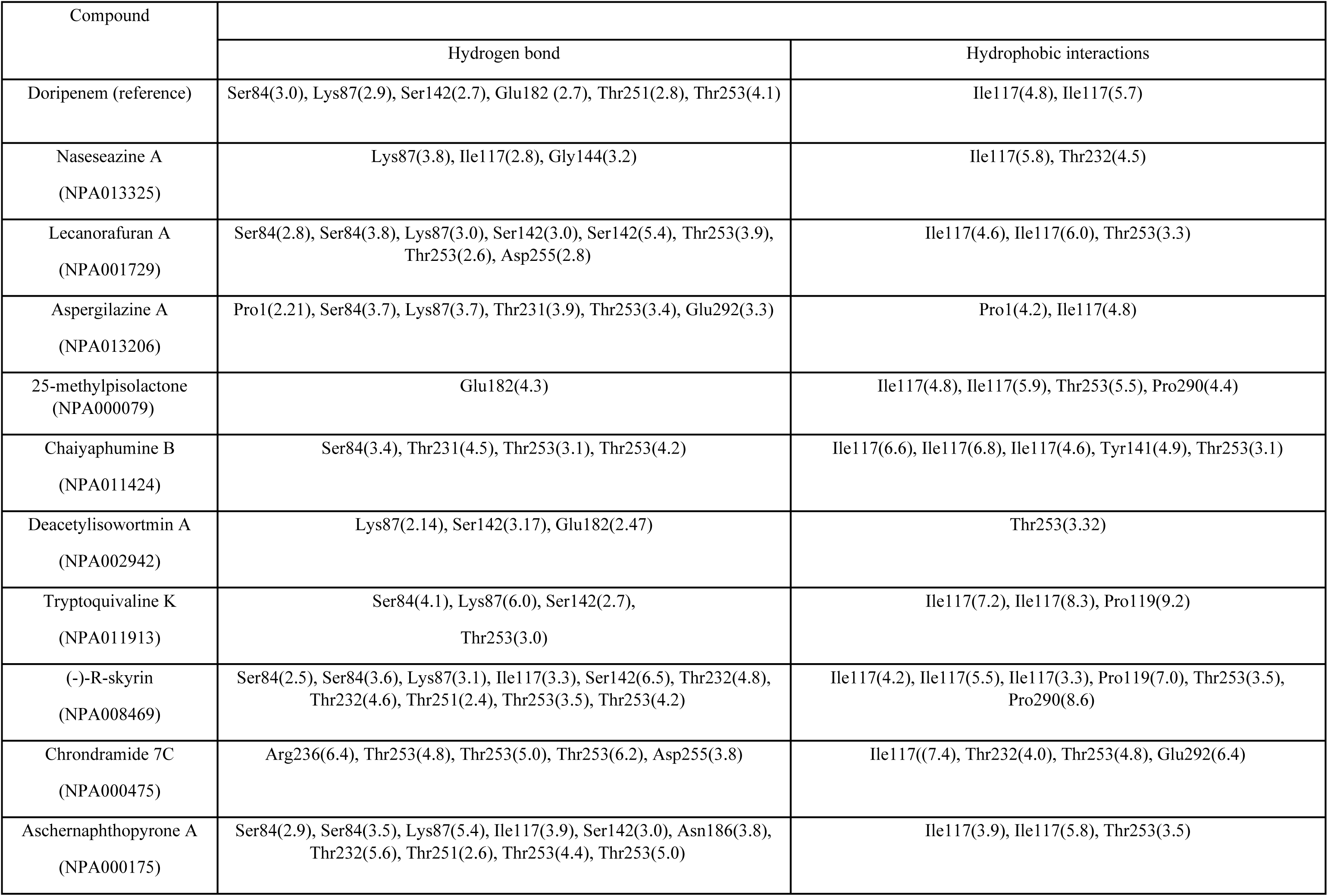
2D interactions of reference and top ligands with *Mycobacterium tuberculosis* beta-lactamase.

Among the top 10 selected compounds’ interactions against BlaC (Fig. 3), NPA013325 with the highest binding affinity showed hydrogen bonds with Lys87, Ile117, and Gly144 residues (Table 3). NPA001729 formed eight hydrogen bonds Ser84, Ser84, Lys87, Ser142, Ser142, Thr253, Thr253, and Asp255. Compound NPA013206 formed six hydrogen bonds with Ser84, Lys87, Thr231, Thr253, and Glu292 residues of BlaC. NPA000079 interacted with several hydrophobic residues and made hydrogen bonds with active site residue Glu182. NPA011424 makes interactions with BlaC via hydrogen bonds Ser84, Thr231, Thr253, Thr253. NPA002942 interacted with Lys87, Ser142, and Glu182 *via* three hydrogen bonds. NPA011913 shows interaction with Ser84, Lys87, Ser142, and Thr253 by making hydrogen bonds with the active site of BlaC. NPA008469 forms hydrogen bonds with Ser84, Ser84, Lys87, Ile117, Ser142, Thr232, Thr232, Thr251, Thr253, Thr253. NPA000475 made interactions via five hydrogen bonds Arg236, Thr253, Thr253, Thr253, and Asp255. NPA000175 formed hydrogen bonds with Ser84, Ser84 Lys87, Ile117, Ser142, Asn186, Thr232, Thr251, Thr253 and Thr253 (Table 3).

From the visualization study of all the hits, we observed that the compounds interact with the active site residues (Ser84, Lys87, Ile117, and Thr253) like the reference molecule Doripenem, suggesting that these compounds can have the ability to inhibit the β-lactamase enzyme. From the results, it was found that the compounds NPA013325, NPA001729, NPA013206, NPA011424, NPA011913, NPA008469, NPA000475 and NPA000175 form additional hydrogen bonds with Gly144, Asn186, Thr231, Thr232, Arg236, Asp255 and Glu292 residues which were not found in the reference (Table 3), suggesting the higher binding affinity of these compounds in the active site of the protein. Further, to substantiate and validate the docking results, multiple docking of each ligand against the protein was performed and the best pose based on the interaction it makes was chosen for further studies.

### 3.4 Pharmacokinetic properties of the selected compounds

Drug likeliness and physicochemical properties of the best ten molecules were calculated using SwissADME(https://www.swissadme.ch/) (Table 4). In general, a compound is considered to be a potential medication if it fits the requirements of Lipinski’s rule of five (Benet et al., 2016). Of the 10 compounds, Lecanorafuran A (NPA001729), Tryptoquivaline K (NPA011913), and Deacetylisowortmin A (NPA002942) had high binding energy (-8.54, - 7.77, -7.79 kcal/mol, respectively), made stable interactions with BlaC (eight, four and three hydrogen bonds respectively) and satisfied Lipinski’s rule of five with zero violations. Therefore, these three compounds were chosen for MD simulation studies to study their stability under the simulated conditions.

**Table 4.**
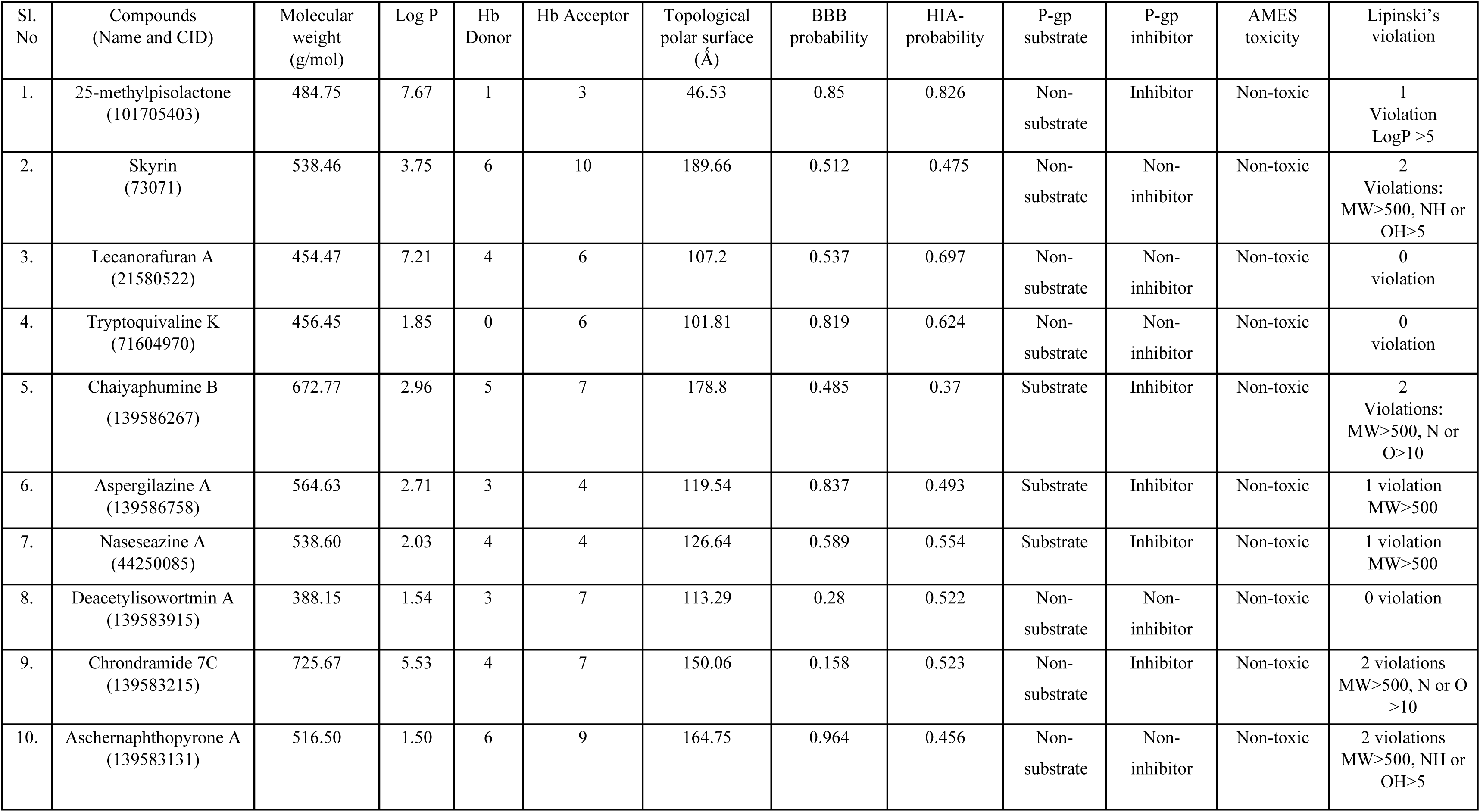
Pharmacodynamic and Pharmacokinetic properties of the top 10 compounds.

Lecanorafuran A is a dibenzofuran identified in the cultured mycobionts of *Lecanora iseana* (Millot et al., 2016) whereas, Tryptoquivaline K is an alkaloid isolated from fungi, especially the genus Neosartorya (Gomes et al., 2014), with both the compounds reported to exhibit antiviral and anti-inflammatory properties. Deacetylisowortmin A was isolated from the endophytic fungus, *Talaromyces wortmanii* (Fu et al., 2016). Surprisingly, all the chemical structures of these compounds lacked the typical beta-lactam ring found in beta-lactamase inhibitors (Table 2).

### 3.5 Molecular Dynamic Simulations

Molecular Dynamic simulations are used to analyze the physical movements of atoms and molecules and to study the conformational changes at the atomic level. MD simulations were performed for 100 ns for the reference (BlaC-Doripenem) as well as the other three selected protein-ligand complexes (BlaC-NPA001729, BlaC-NPA011913, and BlaC-NPA002942) that were shortlisted based on docking binding energy, manual scrutiny for interactions and ADMET analysis.

#### 3.5.1 Conformational stability

Conformational stability was analyzed using RMSD (Root Mean Square Deviation) analysis. RMSD indicates the deviation between the first and other frames generated during simulation. Generally, the lower the RMSD value, the higher the stability of the protein-ligand complex (Aier, n.d.). This assessment offers an understanding of the dynamics and stability of the complexes between the protein and ligands during simulation.

The protein backbone from the crystal structure was selected for the RMSD calculation. The average RMSD values for BlaC-Doripenem, BlaC-NPA001729, BlaC-NPA011913 and BlaC-NPA002942 were 0.38 nm, 0.55 nm, 0.32 nm and 0.53 nm, respectively (Fig. 4a). The average RMSD values shows that BlaC-NPA011913 showed the least RMSD (0.32 nm) value compared to other protein-ligand complexes, including the reference BlaC-Doripenem during the 100 ns simulation. In contrast, BlaC-NPA001729 showed the highest RMSD value (0.55 nm), compared to the other predicted hits, indicating the least stability among the selected complexes. The RMSD of the BlaC-Doripenem reference and BlaC-NPA011913 was stable during the initial phase of the simulation compared to the other two complexes. However, the trajectory of BlaC-NPA011913 started to increase beyond 60 ns due to conformational changes in the rotatable bonds of the ligand. On the contrary, the trajectory of BlaC-NPA002942 showed an unstable trajectory due to the frequent variabilities in the peak during the entire simulation period. To summarize based on the RMSD analysis, the complex formed by the natural product Tryptoquivaline K (NPA011913) with BlaC appears to be more stable and closer to the reference molecule (Doripenem) than the two other hit compounds, Lecanorafuran A (NPA001729) and Deacetylisowortmin A (NPA002942), thereby confirming the stability of the protein-ligand complex (Aier, n.d.).

**Fig. 4.**
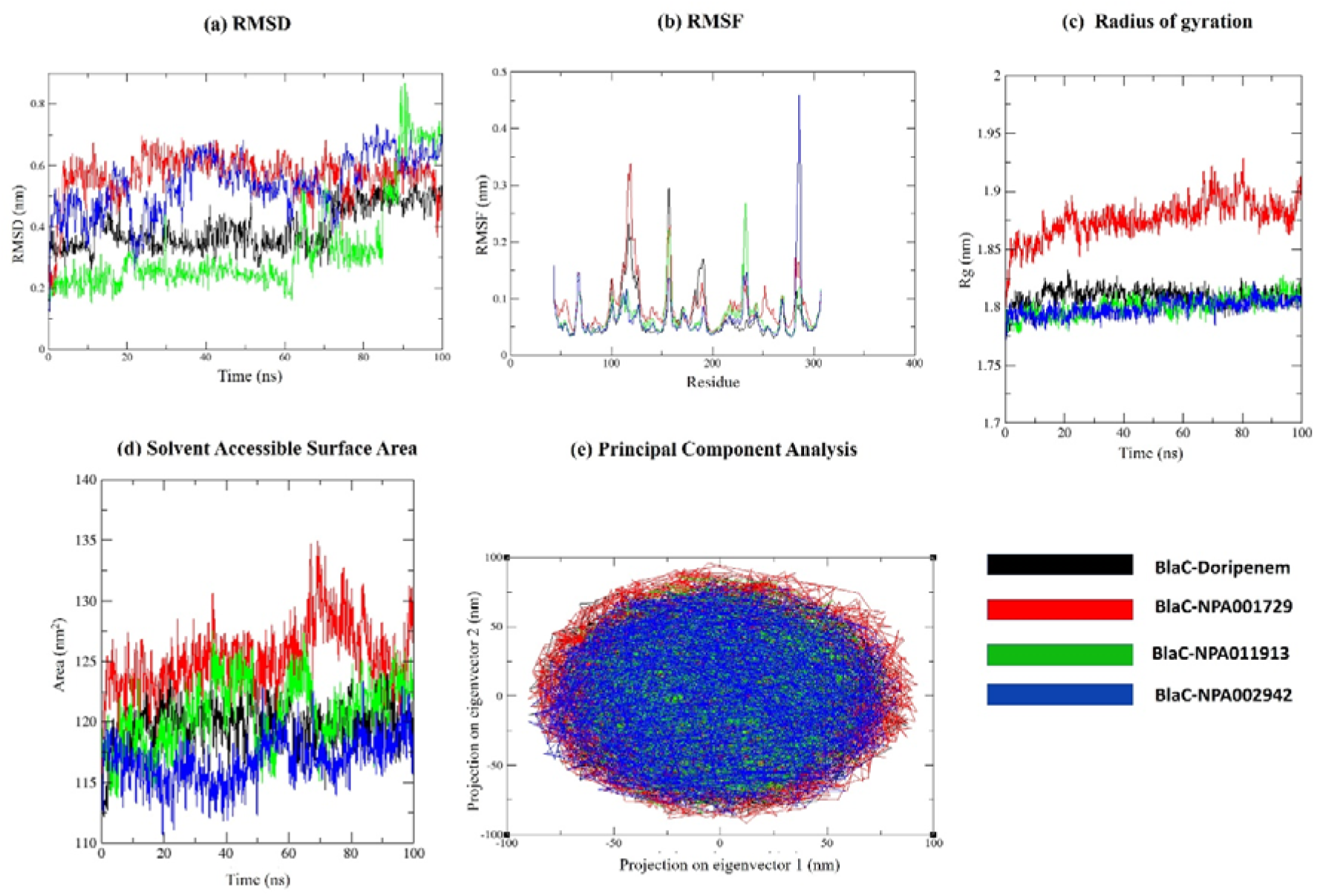
Stability analyses of all the systems. **a** RMSD value for 100 ns **b** RMSF value for the 100 ns trajectory. **c** Radius of gyration vs. the time for 100 ns. **d** SASA value vs. the time for all the systems **e** 2D projection of the first two principal components on the phase space

#### 3.5.2 Fluctuation Analysis

Fluctuation study was performed using RMSF (Root Mean Square Fluctuation) analysis. RMSF represents the fluctuations of residues in a protein upon ligand binding, thereby analyzing the flexibility of different elements in the protein structure. The fluctuations help in understanding atomic-level interactions and structural changes, which helps in drug development as well as protein dynamics. The loosely organized structural components such as turns, loops, and coils generally show higher RMSF values, whereas well-folded structures such as alpha-helix and beta sheets show lower RMSF. Fig. 4b shows the fluctuations at amino acid level upon ligand binding in four different complexes under study. The average RMSF values for BlaC-Doripenem, BlaC-NPA001729, BlaC-NPA011913, and BlaC-NPA002942 were 0.0679 nm, 0.0870 nm, 0.0675 nm, and 0.0697 nm, respectively. In consistent with the RMSD analysis, NPA011913 showed the lowest average RMSF value (0.0675 nm) compared to the reference Doripenem (0.0679 nm) indicating lower fluctuations and higher stability during complex formation with BlaC. On the other hand, the highest average RMSF value was observed for BlaC-NPA001729 (0.087 nm), which embodies higher fluctuation and less stability of the complex. The BlaC-NPA002942 showed a slightly higher average RMSF value (0.0697 nm) compared to the BlaC-DRW complex (0.0679 nm) mainly due to the high fluctuation identified in the C-terminal region (AA 280-290) (Fig. 4b). Thus, the fluctuation analysis of the trajectories indicates that BlaC-NPA011913 is more stable than the reference complex and the other two predicted hits.

#### 3.5.3 Compactness analysis

The Radius of gyration is used for assessing the compactness, rigidity, and folding of a protein. Generally, the lower the Rg value, the higher the stability of the protein while the higher Rg value indicates less compactness in the complex. In this study, the average Rg values were calculated for all four complexes during the 100 ns simulation period (Fig. 4c). The average Rg values for BlaC-Doripenem, BlaC-NPA001729, BlaC-NPA011913, and BlaC-NPA002942 were found to be 1.809 nm, 1.876 nm, 1.799 nm, and 1.805 nm respectively. Surprisingly, the complexes BlaC-NPA011913 and BlaC-NPA002942 showed lesser Rg values (1.799 nm and 1.805 nm) compared to the reference Doripenem complex (1.809 nm) indicating a more stable complex. Furthermore, all complexes except BlaC-NPA001729 (1.876 nm) showed the least fluctuation in Rg values throughout the simulation period suggesting the compactness and stable nature of its complex with BlaC.

#### 3.5.4 Solvent Accessible Surface Area (SASA)

Another parameter to measure the stability of the ligand-protein complex is by analyzing the solvent accessible surface area of the complex. An increased SASA value denotes a less stable complex. Here, the SASA values were calculated for all the complexes (Fig. 4d). The average SASA values BlaC-Doripenem, BlaC-NPA001729, BlaC-NPA011913, and BlaC-NPA002942 complexes were 119.829 nm^2^, 125.406 nm^2^, 116.908 nm^2^ and 116.824 nm^2^, respectively. The SASA analysis of the protein-ligand complexes indicates the lower average SASA values for the BlaC-NPA011913 complex (116.908 nm^2^) and BlaC-NPA002942 complex (116.824 nm^2^) than the control ligand Doripenem complex (119.829 nm^2^) indicating higher stability of the ligand-protein complex, while BlaC-NPA001729 (125.406 nm^2^) shows a higher SASA than the control, indicating the less stable complex formation.

#### 3.5.5 Principle Component Analysis (PCA)

The PCA or essential dynamics calculates correlated motions after ligand binding, to identify the eigenvectors responsible for protein dynamics. PCA analysis highlights the conformational dynamics and stability differences between the reference and selected protein-ligand complexes.

Generally, the first few eigenvectors are important for characterizing the system dynamics. The well-established and less space-occupying cluster represents the stable complex, while the more space-occupying cluster defines the unstable complex. PCA was employed to explore the conformational dynamics of BlaC with Doripenem (reference), NPA001729, NPA011913, and NPA002942. Fig.4e illustrates the conformational sampling by projecting the Cα atoms onto the principal components. The BlaC-NPA011913 and BlaC-NPA002942 complexes showed a very compact cluster with less dispersion, suggesting their higher stability, as they occupy a more confined conformational space. It is worth noting that BlaC-Doripenem and BlaC-NPA001729 did not demonstrate a very compact and dense cluster displaying a dispersed cluster (Fig. 4e). Thus, the 2D PCA suggests that the complexes BlaC-NPA011913 and BlaC-NPA002942 are more stable than the reference BlaC-DRW complex.

#### 3.5.6 Gibbs free energy landscape analysis

Free Energy Landscape (FEL) analysis provides deeper insights into the behaviour of biological systems and conformational dynamics. The Gibbs free energy landscape shows the effect of ligand binding on protein stability. Wherein, the first two principal components (PC1 and PC2) were used to investigate Gibbs free energy landscape. The analysis depicts that the compounds having lower or more significant energy than the reference can follow an energetically more favourable transition from one conformation to another and are thermodynamically favourable.

In figure (5), deep blue represents the lowest energy protein conformation, while red represents the highest energy state protein conformation. The Gibbs energy value ranges from 0 to 5.440 kJ/mol for BlaC-Doripenem, BlaC-NPA011913, and BlaC-NPA002942 complexes, while the Gibbs energy ranges from 0 to 5.820 kJ/mol for the BlaC-NPA001729 complex. The lower free energy values of BlaC-NPA011913 and BlaC-NPA002942 complexes account for the energetically favourable interaction between protein and ligand, signifying a thermodynamically favourable nature of the complex similar to the reference.

**Fig. 5.**
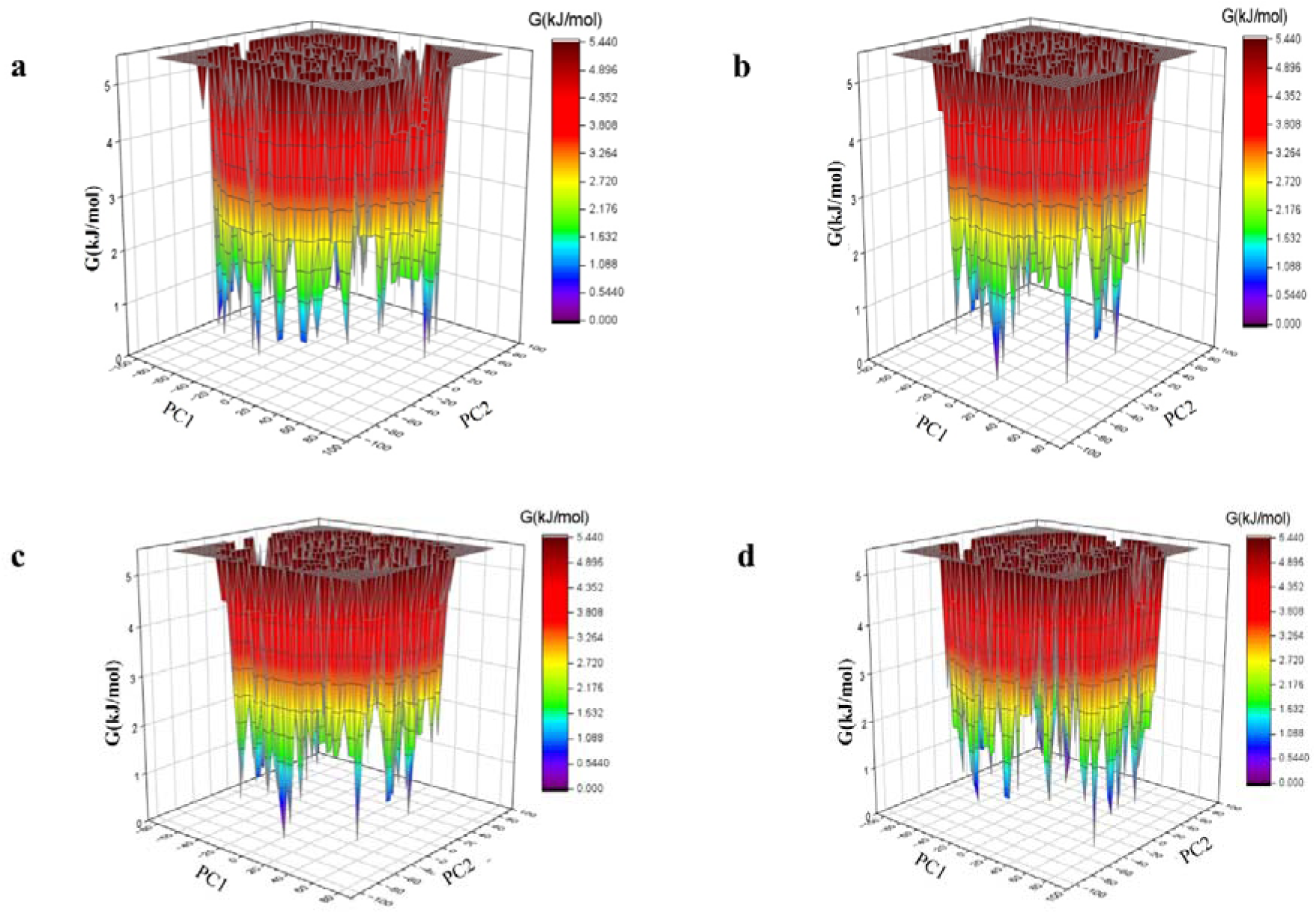
Gibbs free energy landscape. **a** BlaC–DRW, **b** BlaC-NPA001729, **c** BlaC-NPA011913, **d** BlaC-NPA002942

### 3.6 Molecular Mechanics/Poisson-Boltzmann Surface Area (MMPBSA) analysis

The binding free energy calculation provides insights into the stability of biomolecular interactions of the complexes. Various energetic terms such as van der Waals energy, electrostatic energy, polar solvation energy, and binding free energy were calculated and shown in Table 5. The binding free energy for BlaC-Doripenem, BlaC-NPA001729, BlaC-NPA011913, and BlaC-NPA002942 was calculated to be –64.39 kJ/mol, +1807.15 kJ/mol, - 134.39 kJ/mol and -33.47 kJ/mol respectively. Consistent with all our prior analyses, BlaC-NPA011913 (-134.39 kJ/mol) has a total binding energy lower than the reference compound Doripenem (-64.39 kJ/mol), indicating its higher stability and improved binding affinity compared to the reference compound. While BlaC-NPA002942 and BlaC-NPA001729 showed a higher binding energy than the reference complex indicating a weaker protein-ligand complex formation.

**Table 5.**
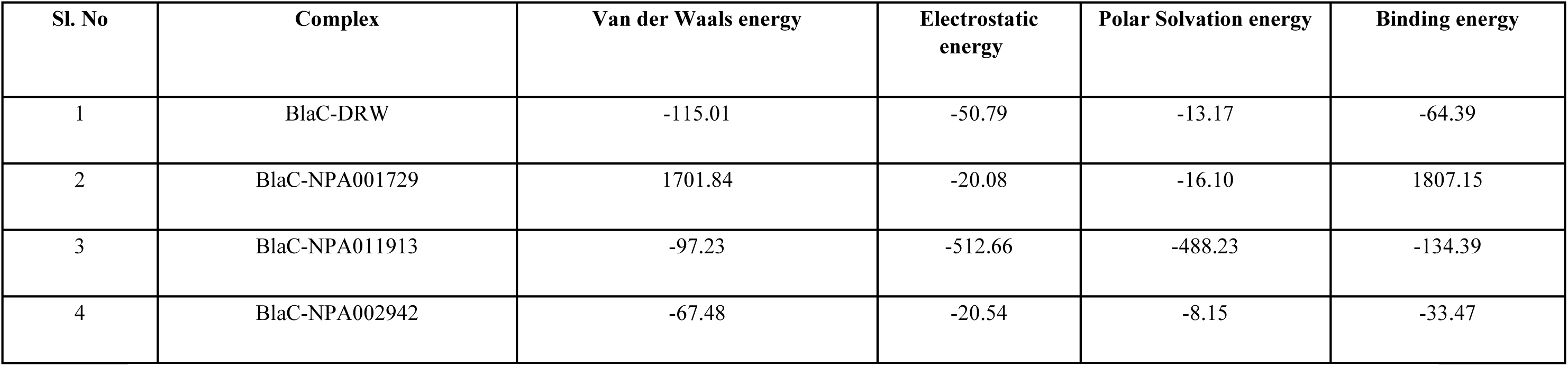
Electrostatic, Polar solvation, and Binding energies in kJ/mol for the control compound (DRW) and predicted hits.

| Sl. No | Complex | Van der Waals energy | Electrostatic energy | Polar Solvation energy | Binding energy |
| --- | --- | --- | --- | --- | --- |
| 1 | BlaC-DRW | -115.01 | -50.79 | -13.17 | -64.39 |
| 2 | BlaC-NPA001729 | 1701.84 | -20.08 | -16.10 | 1807.15 |
| 3 | BlaC-NPA011913 | -97.23 | -512.66 | -488.23 | -134.39 |
| 4 | BlaC-NPA002942 | -67.48 | -20.54 | -8.15 | -33.47 |

Overall, based on the results obtained through various *insilico* analyses and validations, the compound NPA011913 displayed strong binding affinity and stability with BlaC protein compared to that of the other selected compounds. The identified compound showed strong potential as a BlaC inhibitor, with binding scores and interaction patterns similar to known inhibitors. Further, *invitro* and *invivo* studies are warranted in this regard.

## 4. CONCLUSION

Multidrug resistance exhibited by *Mycobacterium tuberculosis* has triggered the need for the development of new anti-TB drugs. In this study, natural compounds were subjected to molecular docking and MD Simulation studies against the β-lactamase (BlaC) of *Mycobacterium tuberculosis*. To understand the efficacy of these compounds, BlaC complexed with a known inhibitor, doripenem, was considered as standard. Based on virtual screening and insilico docking analysis, it was observed that the natural compounds Lecanorafuran A, Tryptoquivaline K and Deacetylisowortmin A showed significant binding affinities with the target protein. Among the top 3 compounds, Lecanorafuran A exhibited considerably higher binding affinity among the three compounds. However, the MD Simulation analysis illustrating the stability of the interactions observed in the docked complexes through RMSD, RMSF, Rg and energy profiles, showed that the compound Tryptoquivaline K was found to form a stable complex with BlaC than the other two selected compounds. The findings of this study highlight the potential of natural compounds with no beta-lactam ring as a replacement to the chemically synthesized inhibitors with beta-lactam ring in targeting the BlaC protein of *M. tuberculosis*. However, it is important to note that these findings are based on *insilico* techniques. Further experimental validations are required to confirm the efficacy of Tryptoquivaline K as an inhibitor of *Mycobacterium tuberculosis* β-lactamase. In addition, these compounds should be tested for their pharmacokinetic properties and other potential undesired targets. Nevertheless, the results of this study provide valuable information on the potential of natural compounds from various sources as inhibitors of BlaC, suggesting an *in vitro* study of these compounds as promising drug candidates.

## ACKNOWLEDGEMENTS

We sincerely thank the Vellore Institute of Technology, Vellore for their support.

## AUTHOR’S CONTRIBUTION

I.D. conceptualized the study, and S.S. performed the computational study, S.S., B.J.L., and L.S.S analysed the results, S.S., S.C.P., and N.E.R. prepared the manuscript draft with figures. I.D. reviewed and interpreted the obtained results. A.P.S. and T.R. reviewed the manuscript, provided critical feedbacks on analysis performed and improved the manuscript. I.D. approved the final manuscript.

## DECLARATION

### Ethics Approval

This article is an *insilico* work-based research article and thus studies with either humans or animals were not performed for this research by any of the authors.

